# Modeling of Yield Losses and Risk Analysis of Fungicide Profitability for Managing Fusarium Head Blight in Brazilian Spring Wheat

**DOI:** 10.1101/608489

**Authors:** Maíra Rodrigues Duffeck, Kaique dos Santos Alves, Franklin Jackson Machado, Paul David Esker, Emerson Medeiros Del Ponte

## Abstract

Fusarium head blight (FHB), caused by the *Fusarium graminearum* species complex, is a serious disease of wheat in Brazil. A review of literature on fungicide efficacy for field trials evaluated in Brazil was conducted to obtain FHB-yield data and explore their relationship. Thirty-seven studies (9 years and 11 locations) met the criteria for inclusion (FHB index ≥ 5% and max-min range ≥ 4 percent points [p.p.]). Studies were group into two production situations: low (*Yl* ≤ 3,631 kg ha^−1^) or high (*Yh* > 3,631 kg ha^−1^) yield, based on the median of maximum yields across trials. Population-average intercepts, but not the slopes, from fitting a random-coefficients model, differed significantly between *Yl* (2,883.6 kg ha^−1^) and *Yh* (4,419.5 kg ha^−1^). The calculated damage coefficient was 1.05 %^−1^ and 1.60 %^−1^ for *Yh* and *Yl*, respectively. A crop model simulated attainable wheat yields for 10 planting dates within each year during a 28-year period, including prior (1980-1989) and after (1990-2007) FHB resurgence. Simulated losses using disease predictions to penalize yield were in general agreement in magnitude with literature reports, for both periods. Economic analysis for scenarios of variable fungicide costs and wheat prices, and one versus two sprays of tebuconazole, showed that the probability of not-offsetting the costs was higher (> 0.75) prior to FHB resurgence than after the 1990. Our approach may be useful for designing of longlasting, yet profitable, contingency tactics to management FHB in wheat. Currently one spray of triazole fungicide during flowering is more likely a profitable decision than applying two sprays, for which there is greater uncertainty.

## Introduction

Fusarium head blight (FHB) is a flower-infecting fungal disease of wheat and barley that directly and negatively impacts yield and contaminates grains with mycotoxins produced by the fungus (Goswami and Kistler 2004; McMullen et al. 1997). FHB has been considered a resurgent problem worldwide since the early 1990s with economic losses estimated around $2.7 billion in the United States over a two decade period (McMullen et al. 2012). Similar patterns have been reported in wheat and barley regions in South America where the frequency of FHB epidemics increased during the 1990s compared to the previous two decades (Fernandes 1997; Kikot et al. 2011; Moschini and Fortugno 1996). Two hypotheses for this resurgence include the wider adoption of conservation tillage practices, as well as the increased use of corn-wheat rotations in North America and Europe (Landschoot et al. 2013; Schaafsma et al. 2005). In Brazil, modeling work also showed a significant contribution of climate variability to the resurgence of FHB based on analysis of a 50-year weather dataset for a location in southern Brazil (Del Ponte et al. 2009).

The occurrence and intensity of FHB epidemics are driven by the amount of airborne inoculum (primarily ascospores), which may originate from both within and outside the field, combined with humid conditions during and post anthesis period that favor infection of the host (McMullen et al. 2012). Disease control is best achieved through the use of integrated practices including cultural measures aimed at reducing local inoculum, the use of moderately resistant cultivars and applying fungicide around anthesis (Wegulo et al. 2011). Although chemical fungicides, mainly of the triazole chemistry, have been recommended for controlling FHB, the efficacy levels are generally modest and influenced the type of active ingredient, wheat cultivar, number of sprays and application technology (Edwards and Godley 2010; Machado et al. 2017; Paul et al. 2008; Willyerd et al. 2012).

Coinciding with the resurgence of FHB, research since the mid-1990s has focused on evaluating a range of control methods, including host resistance and fungicide sprays, all with the explicit goal to reduce mycotoxin contamination and protect wheat yields. Thus, these efforts have generated a large amount of epidemiological data (Kazan et al. 2012; McMullen et al. 2012; Shah et al. 2018). These large data sets have enabled the use of meta-analytic approaches to study the effects of fungicides on disease control, yield increase and mycotoxin reduction, as well as to explore empirical relationships between FHB index and wheat yield or mycotoxin levels (Madden and Paul 2009; Salgado et al. 2011, 2014, 2015). Meta-analysis has become standard for studying empirical relationships between disease and yield data for other diseases like soybean rust (Dalla Lana et al. 2015), soybean white mold (Lehner et al. 2017) and soybean target spot (Edwards Molina et al. 2018).

The majority of studies on the relationship between FHB intensity and variables like yield or mycotoxin have been conducted in the temperate regions in North American cropping conditions (Madden and Paul 2009; Salgado et al. 2011), which differ from subtropics of South America in both abiotic and biotic factors, as well as management practices (Del Ponte et al. 2015; Spolti et al. 2014). In North America, for example, the relationship between FHB index and yield was quantified using field trials from multiple location-years to estimate damage coefficients (percent reduction in yield per percent point increase in FHB index) for winter wheat (Madden and Paul 2009; Salgado et al. 2015) and spring wheat (Madden and Paul 2009).

In Brazil, previous efforts to estimate yield losses due to FHB were based on destructive sampling of wheat heads in replicated non-treated plots of field experiments to estimate attainable (non-diseased spike/samples weight) and actual yield (sample weight) based on absolute and relative yield losses. Using this method, mean yield losses were estimated to be higher for the period 1999-2010 compared to the 1984-1994 period (Panisson et al. 2003; Reis et al. 1996; Reis and Carmona 2013). Since the early 2010s, significant yield reductions due to FHB have been reported in non-treated plots from a network of uniform fungicide trials (UFTs) (Santana et al. 2012, 2014, 2016a, b, c). A recent meta-analysis of the effect of fungicides on FHB control and yield using data from several sources, including the UFTs data from Brazil, explored yield responses and economic benefits from using fungicides. Results suggested that one spray of triazole fungicide may suffice and be economic to manage the disease (Machado et al. 2017).

While several papers have highlighted the impact of FHB on wheat in Brazil, the heterogeneity of the functional relationship between FHB intensity and wheat yield has not been explored. This approach could be a useful non-destructive method for predicting FHB index and improving the risk assessment of economic losses due to FHB. Thus, the objective of this study was to model the relationship between a single (critical-point) assessment of FHB index and wheat yield data obtained from independent studies in order to obtain estimates of damage coefficients in a manner similar to other previously published studies (see for example, Edwards Molina et al. 2018; Lehner et al. 2017; Madden and Paul 2009; Salgado et al 2015). To reconstruct yield losses curves in a 28-year time series, covering the period before and after FHB resurgence, the damage coefficient was used to penalize attainable yields simulated by a crop model based on predictions of FHB index by a disease model. Furthermore, the risk of not-offsetting on the cost of applying fungicides (one or two-spray treatment) was calculated for a range of wheat prices and fungicide costs for each year.

## Material and Methods

### Data source, description and criteria for inclusion

A systematic review of peer- and non-peer-reviewed articles/reports published after the year 2000 was conducted to identify studies reporting wheat yield and FHB intensity data in Brazil. In total, 29 publications reported results from trials conducted at 23 locations in southern Brazil spanning a 16-year period (2000 to 2016). Altogether, these locations comprise 90% of the total wheat production in Brazil (CONAB 2018) (Fig. S1). Portion of these data has been used in a previous study on the effect of fungicides on disease and yield (Machado et al. 2017). In brief, the studies were conducted following a standard protocol and regional recommendations of agronomic practices. Fusarium head blight disease development was natural in all trials. The number of treatments evaluated in each trial ranged from 5 to 13 (average of 8.5 treatments/trials). All trials included a non-treated control. Fungicides were typically applied once at flowering (Zadoks scale 60 – 64) or twice, being the first at flowering and the second 10 days later. The specific fungicide treatments, and respective application rates, varied among trials (Machado et al. 2017).

The disease response variable used in our study, FHB index (%), which represents the proportion of diseased spikelets in the sample of spikes (Paul et al 2005, 2007). This index was assessed during soft dough stage (Zadoks scale 85), prior to senescence. Plots were harvested and the mean yield (kg ha^−1^) adjusted to 13% grain moisture. Data from all trials were inspected closely for both FHB index and yield from which two criteria were defined to select trials for inclusion in the analysis: a) FHB index in the trial should be at least 5%, to ensure that yield reductions were due to FHB, and b) a minimum range of four percentage points between the minimum and maximum FHB index should occur in order to more reliably estimate the slopes (Madden and Paul 2009). Twenty-seven trials did not meet these minimum criteria (primarily due to FHB index < 5%) and were excluded from subsequent analyses. Influence analysis was also performed from which five influential trials were excluded (data not shown). Therefore, data were available for 37 trials reported in 18 studies (Table S1).

### Effect sizes and meta-analytic modeling

Intercepts (*β*_*0*_) and slopes (*β*_*1*_) (using ordinary least square regression modeling) were estimated for the relationship between FHB index and wheat yield at the trial level (Dalla Lana et al. 2015; Madden and Paul 2009). The distribution of the linear coefficients estimated independently for each trial was summarized by calculation of the interdecile (ID) range (Lehner et al. 2017; Madden and Paul 2009).

A multilevel (random coefficients) model was fitted to the yield-FHB index relationship and the population-average intercept and slope were estimated as described elsewhere, assuming a linear relationship between FHB index and yield (Lehner et al. 2017; Madden and Paul 2009). The predicted study-specific intercepts 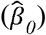 and slopes 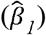 were estimated from the best linear unbiased predictors (BLUPs) using maximum likelihood, together with the population-average parameters. Models were fitted separately for trials based on two production situations that were defined based on maximum yield achieved in the trial: high yield (*Yh*) or low yield (*Yl*). Yields in the low-yielding category of trials were assumed to be due to unknown (unmeasured) abiotic (low soil fertility, drought) or biotic stresses (foliar diseases, pests, etc). The classification of individual trials was based on the median of the maximum yield values across trials (3,631 kg ha^−1^).

The *lmer* function of the *lme4* package of R was used to estimate the mean effect based on the between-study variance and within-study variance and also predict the study-specific intercept (*β*_*0*_) and slope (*β*_*1*_) coefficients (Lehner et al. 2017). Yield category was tested in the mixed effects model as a moderator variable in order to account for the heterogeneity in the intercept and/or slope of the yield-FHB index relationship. Wald-type tests were performed to determine if the inclusion of the covariate in the model significantly affected the model coefficients. A damage coefficient (*Dc*), which is commonly reported in relative terms (percent reduction in yield in our study), was calculated in order to compare with other studies. The *Dc* for the low-yield and high-yield production situations was calculated by dividing the estimated population-average slope 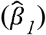 (kg ha^−1^%^−1^) by the estimated population-average intercept 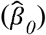 (kg ha^−1^) times 100 (Lehner et al. 2017; Madden and Paul 2009).

### Simulation of yield losses in the historical time series

The calculated damage coefficient (*Dc*) was used to predict yield losses by linking a wheat growth and yield simulation model to a FHB risk infection simulation model (Del Ponte et al 2005; Del Ponte et al. 2009). This was done over a three decade period that encompassed two different FHB periods which we called “prior” and “after” FHB resurgence in Brazil. First, a crop simulation model (cropsim-wheat) calibrated for a spring wheat cultivar (BRS Guamirim) and soil profiles of conditions at Passo Fundo (28°15’00”S and 52º25’12”W) was used to simulate attainable yield (Lazzaretti et al. 2015), or yield constrained only by factors like radiation, temperature, crop phenology and physiology, as well as water and soil nutrients (Savary et al. 2018), during ten planting dates over a 28-year period (1980 to 2007). The 10 yearly runs were performed from sowing dates ranging from 1 June to 30 June (spaced three days apart), which is the recommended sowing period for Passo Fundo. Starting on each heading date predicted by the wheat model, the FHB model predicted daily infection risks, based on daily weather conditions (Del Ponte et al. 2015). The daily risks were accumulated during flowering period and further used to predict FHB severity index based on a linear regression model (Del Ponte, et al. 2005). A similar approach was used to simulate FHB index (30 planting dates after June 1^st^) for a 50-year dataset (Del Ponte et al. 2009). The attainable yield predicted by the crop model was penalized by the *Dc* using the predicted FHB index for each sowing date. For example, for the sowing date of 6 June 1990, simulated yield and FHB severity were 3645.38 kg ha^−1^ and 19.5%, respectively, and for this situation FHB reduces yield by 20.5% (or 747.17 kg ha^−1^). In total, 280 simulations (10 planting dates in 28 seasons) of yield reductions were performed.

To test for significance of the trends in yield loss predictions in the time series, as well as to test the effect of two sowing periods (sowing dates before or after June 15^th^), a generalized additive model (GAM) (Wood 2017) was fitted to the data using the *gam* function of *mgcv* R package (Wood 2017). The smooth terms estimated for each sowing period were compared at 5% significance.

### Risk analysis of not-offsetting control costs

An economic analysis to quantify the probability of not-offsetting control costs was conducted, aiming to verify when the cost of the fungicide was not covered by an expected relative increase in yield (kg ha^−1^). This analysis was conducted in several steps, with the probability values being obtained through simulation approach. The first step was to extract the estimates of control efficacy for tebuconazole fungicide used in FHB management in Brazil, which were already summarized by Machado et al. (2017). Therefore, the obtained estimates of control efficacy (*C*) for tebuconazole, either applied once (1□) or twice (2□), were used in our simulations. For this, we assumed *C* to follow a uniform distribution given by:

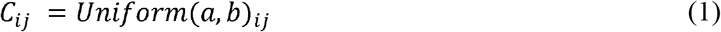

being *i* the planting date and *j* the year, totaling 280 simulations of control efficacy (10 planting dates in June) for one (1□) and twice (2□) fungicide application; *a* and *b* represent the lower (*CI*_*L*_) and upper (*CI*_*U*_) limits of the 95% confidence interval around *C*. The *CIs* ranged from 46.9% to 67.5% for one (1□) fungicide application, and from 44.3% and 60.7% for two fungicide applications (Machado et al. 2017).

The simulated *C* values were used to calculate FHB severity index, and respective yield response, in the fungicide-treated plots. The mean yield difference (*D*) for one spray was given by the difference in yield between fungicide-protected (*Yield*_*f*_) and no fungicide (*Yield*_*nf*_) plots (Machado et al. 2017; Madden et al 2016). The *D* from using two sprays was calculated by adding a simulated (normally distributed) mean yield gain from using a second spray of tebuconazole, as determined in the previous meta-analysis study (Machado et al. 2017) (Equation 2).

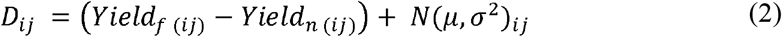

being *i* the planting date and *j* the year, *μ* is 101.6 kg ha^−1^, representing the mean yield difference between the 1□ and 2□ fungicide application (Machado et al. 2017), and *σ* is 43.8, that was calculated based on the mean of the lower (*CI*_*L*_) and upper (*CI*_*U*_) limits of the 95% confidence interval around *D* for one spray. Lastly, the between-year variance 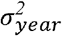 was estimated based on the ten values of *D* within each year, or across planting dates.

The estimates of *D* and 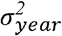 were used to calculate the risk (probability) of not-offsetting fungicide application cost (*P*_*loss*)_, taking both fungicide application cost (*F*_*c*_ = fungicide plus application costs) and the average wheat price (*W*_*p*_) by year. This probability is the cumulative standard-normal function given by:

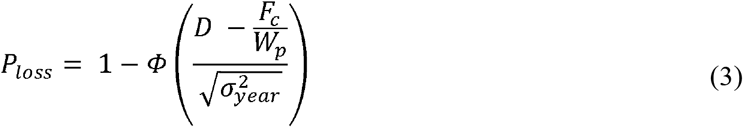

where *Φ* is the cumulative standard-normal function (Machado et al. 2017; Paul et al. 2011).

Different scenarios of and *W*_*p*_ were *F*_*c*_ simulated using average prices for *F*_*c*_ ranging from $5,00 to $35,00 ha^−1^, and average prices of *W*_*p*_ ranging from $100,00 to $250,00 ha^−1^. The minimum and maximum *F*_*c*_ were defined based on previous studies to in Brazilian conditions (Machado et al. 2017; Panisson et al 2003; Reis et al. 1996). Likewise, the range of prices for *W*_*p*_ were defined based on published statistics for southern Brazil (Brum et al. 2005; CEPEA, 2018). Tile (rectangle) plots were produced with classes of risk (*P*_*loss*_) (*p* < *25%,25* ≤ *p* < *50%,50* ≤ *p* < *75%,p* ≥ *75%*), represented by different colors, for each *F*_*c*_ × *W*_*p*_ combinations for each of the three periods (1980-1989, 1990-1999, 2000-2007). A simplified diagram depicting the steps of our modeling approach is presented in Figure 1.

**Figure 1.**
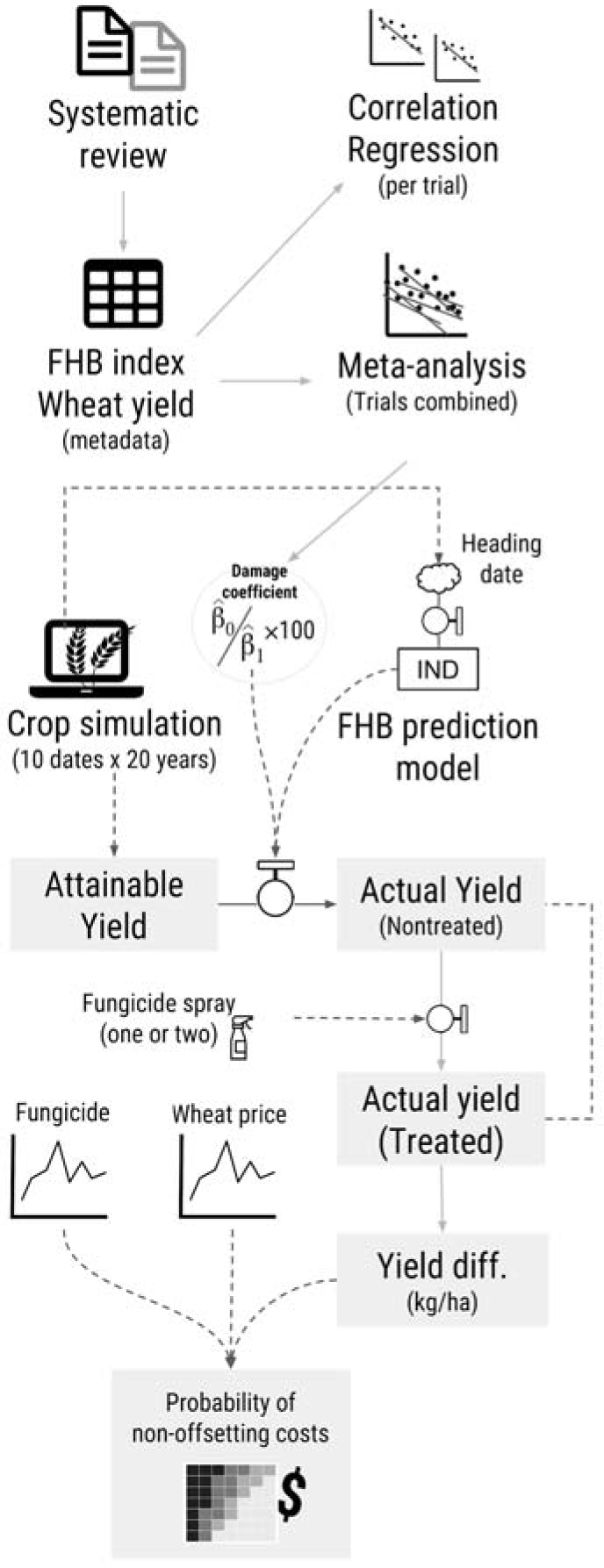
Simplified diagram of the steps taken to modeling yield losses caused by Fusarium head blight in spring wheat in Brazil and to quantify the economic impact of fungicide application.

### Data availability and reproducibility

All data gathered for this study as well as the fully annotated computational codes, prepared in RMarkdown, were organized as a research compendium publicly available at https://osf.io/3h9ye/. A website was generated to facilitate visualization of the commented scripts (https://emdelponte.github.io/paper-FHB-yield-loss/) All analyses were conducted in R statistical software (R Core Team, 2013).

## Results

### Yield-severity relationship

#### Simple linear regression model

Across all studies, FHB index in the non-treated plots ranged from 5 to 89.3% (mean = 23.6%). In 19 (51.3%) trials FHB index was greater than 10%, which is considered a threshold for epidemics of concern (Shah et al. 2013). Maximum yield ranged from 1,460 to 5,692 kg ha^−1^ (mean = 3,676 kg ha^−1^). Intercepts and slopes varied considerably within each yield group. For *Yl*, intercepts ranged from 1,708 to 4,299 kg ha^−1^, with an overall mean of 2,977 kg ha^−1^ and slopes ranged from 43.5 to −381.9 kg ha^−1^, with overall mean of −71.8 kg ha^−1^ (Fig. 2A). For the *Yh* crop situation, intercepts ranged from 3,000 to 6,097 kg ha^−1^, with an overall mean of 4,473 kg ha^−1^ (Table S2) and slopes ranged from 52.9 to −220.2 kg ha^−1^, with overall mean of −60.9 kg ha^−1^ (Fig. 2A). All but two slopes (*β*_*1*_) were negative, one in each yield group, indicating that yield generally decreased as FHB index increased.

#### Random coefficients model

The multilevel model that considers the random effect of individual trials on both the intercept and slope provided a better fit to the data, compared with slope or intercept-only models (data not shown), across the 37 trials. Mixed model-derived intercept and slope values varied among individual studies (Table S2). Study-specific intercepts 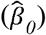 ranged from 1,620 to 5,786 kg ha^−1^(ID = 2,470.6 kg ha^−1^) and study-specific slope 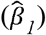 values ranged from 4.5 to 112.5 kg ha^−1^ %^−1^ (ID = 76.8). The estimates of population-average of the intercept and slope were 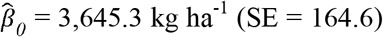 and 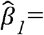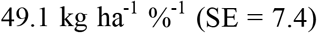, respectively. Both parameters were statistically different from 0 (*P* < 0.001).

The best model based on likelihood ratio test (*P* < 0.001) and lowest AIC (4,595) included baseline yield as a covariate. Based on a Wald-type test, *ς* (effect of baseline yield on the slope) did not differ from 0 (*P* > 0.80); suggesting that the slope was not affected by baseline yield. However, as expected, *ξ* (effect of baseline yield on the intercept) differed from zero (*P* < 0.01). The study-specific parameters estimated were: 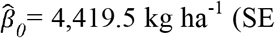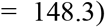, 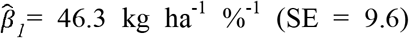, and 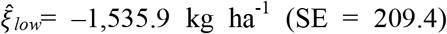. Population-average predictions were then split into two regression equations to account for the effects of the covariate on the intercept. In addition, *ξ* was a dummy variable of value of zero for high yield (i.e., 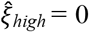). Because of the negative value for 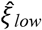, wheat yield was, on average, 1,535.97 kg ha^−1^ less than in the high-yield trials. The population-average predictions of yield are given by:

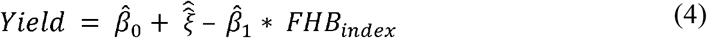

which gives the following equations for high and low wheat yield:

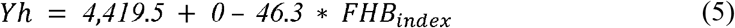

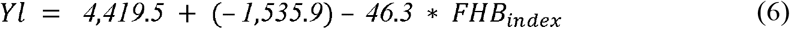

The population-average (and respective 95% *CI*_*s*_) predictions for the inclusion of the covariate baseline yield in the model are shown in Figure 2B. Because the estimated slope did not differ between low and high yield class, a reduction of 1,000 kg ha^−1^ would occur at an FHB index of 21.6 % for both yield classes. The damage coefficient predictions for high-yield and low-yield conditions were 1.05%^−1^ and 1.6%^−1^, respectively.

**Figure 2.**
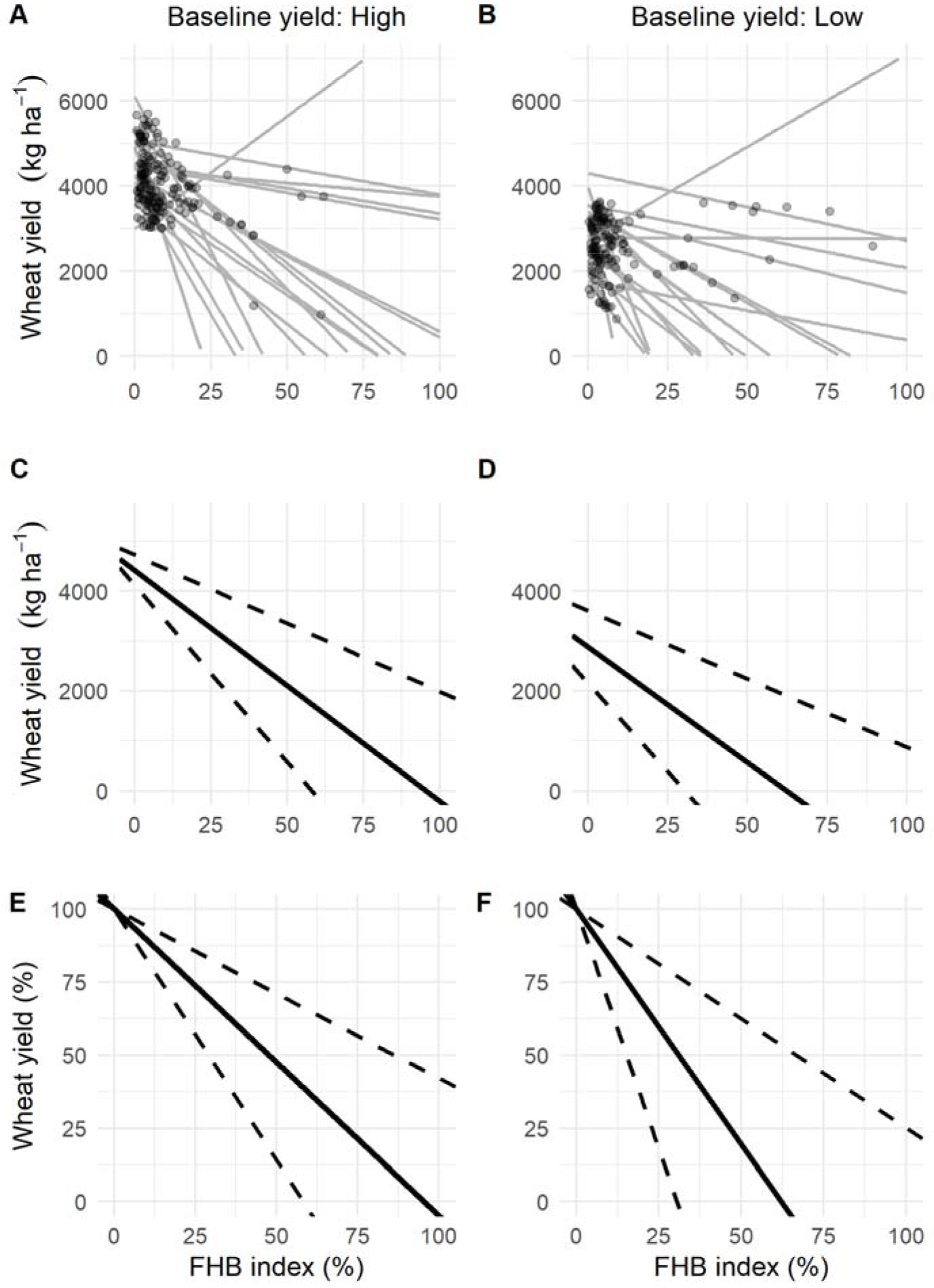
Results for the fit of a simple linear model to wheat yield (kg ha^−1^) and FHB index (%) for both crop situations of **A**, high yield or **B**, low yield. Results for the fit of a random-coefficient model to wheat yield (kg ha^−1^) and FHB index (%) with the population-average predictions (thick solid black) of absolute (kg ha^−1^) and relative yield (%) and respective 95% confidence interval (thick dashed black) for **C** and **E**, high yield or **D** and **F**, low yield.

### Model predictions of yield losses

Attainable yield simulations based on the crop model were consistently above the baseline yield (3,631 kg ha^−1^). Yield estimates were then, penalized by the *Dc*_*h*_ (1.05% pp^−1^) representing high-yielding situation. Overall, relative yield reductions varied significantly both intra and inter-annually (Fig. 3A). For the pre-FHB resurgence period (1980-1989), yield losses ranged from 1.0% to 12.6% (mean = 5.8%). For the two subsequent periods, 1990-1999 and 2000-2007, yield losses ranged from 3.1% to 22.6% (mean = 10.3%) and from 3.2% to 25.8% (mean = 10.2%), respectively (Fig. 3A). A significant upward trend was detected in the time series (*P* < 0.01). However, differences between the smooth terms of the two planting date periods within a season were not significant (*P* > 0.05) (Fig. 3B).

**Figure 3.**
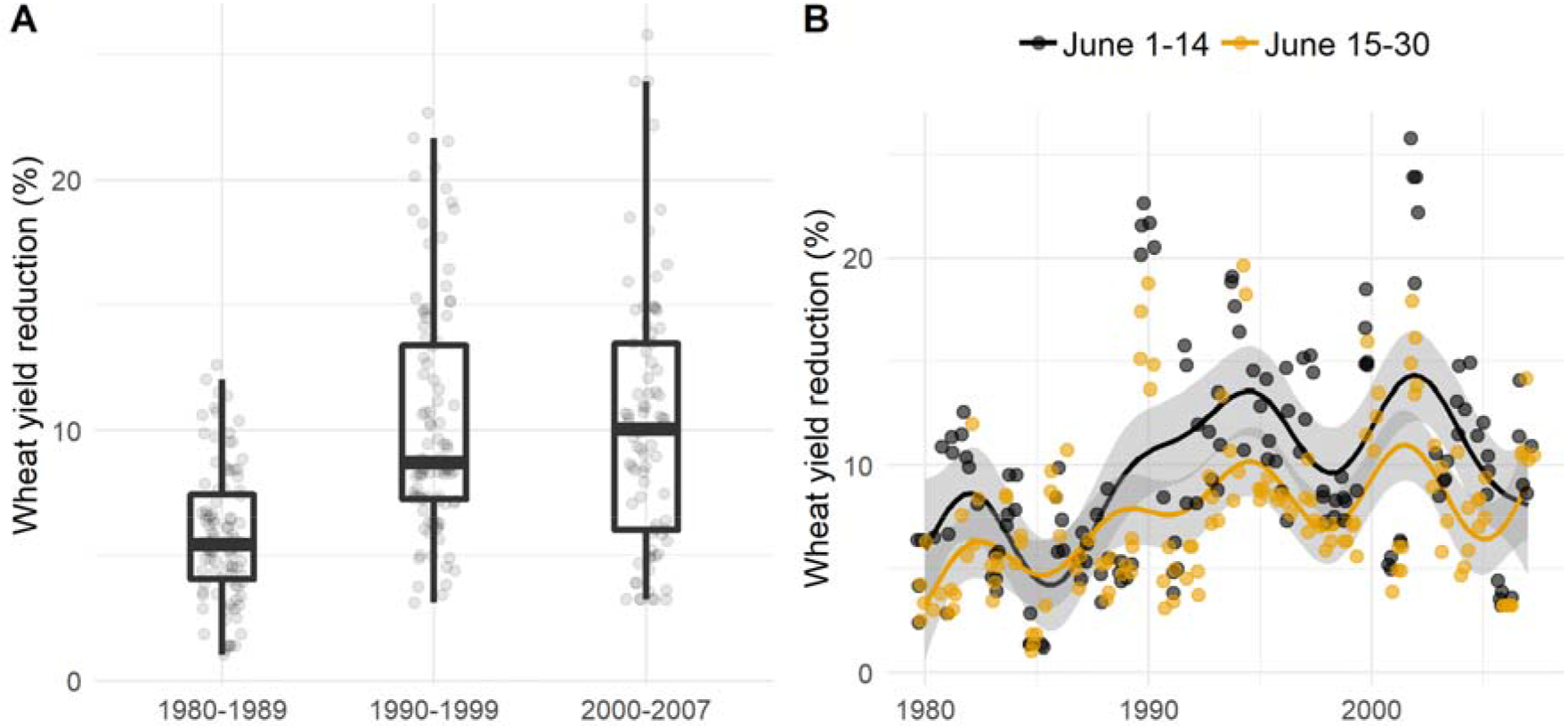
Relative yield loss estimated by a phenology-based model in a 28-year period (1980 to 2007) for Passo Fundo, RS, Brazil. Box-plots in **A** represent the variability of the relative losses within three time periods. Relative yield loss in **B** represents the estimated values encompassing the sowing dates before and after 15 June, with the fitted smoothing lines of the generalized additive model for both sowing periods.

### Probability of not-offsetting fungicide costs

When the three periods were compared, the probability of loss (*P*_*loss*_) was greater during the pre-FHB resurgence period (1980-1989) than for the two subsequent periods, regardless of the number of sprays (Fig. 4A, B). The *P*_*loss*_ was more affected by fungicide cost and wheat prices than the number of fungicide sprays. For example, for a fungicide spray costing $20 U.S./ha, and wheat price of $175 U.S./ton, *P*_*loss*_ was 59%, 26%, and 35% for 1980-1989, 1990-1999, and 2000-2007, respectively. When using two sprays, the values were 59%, 33%, and 34%, for 1980-1989, 1990-1999, and 2000-2007, respectively. This is also clear in the yearly simulations *P*_*loss*_ of for more likely benefit-cost ratios (wheat price / fungicide cost in US$) where the risk of not offsetting the costs tended to decrease after the early 1990s and remained lower for scenarios of one than two sprays (Fig. 5).

**Figure 4.**
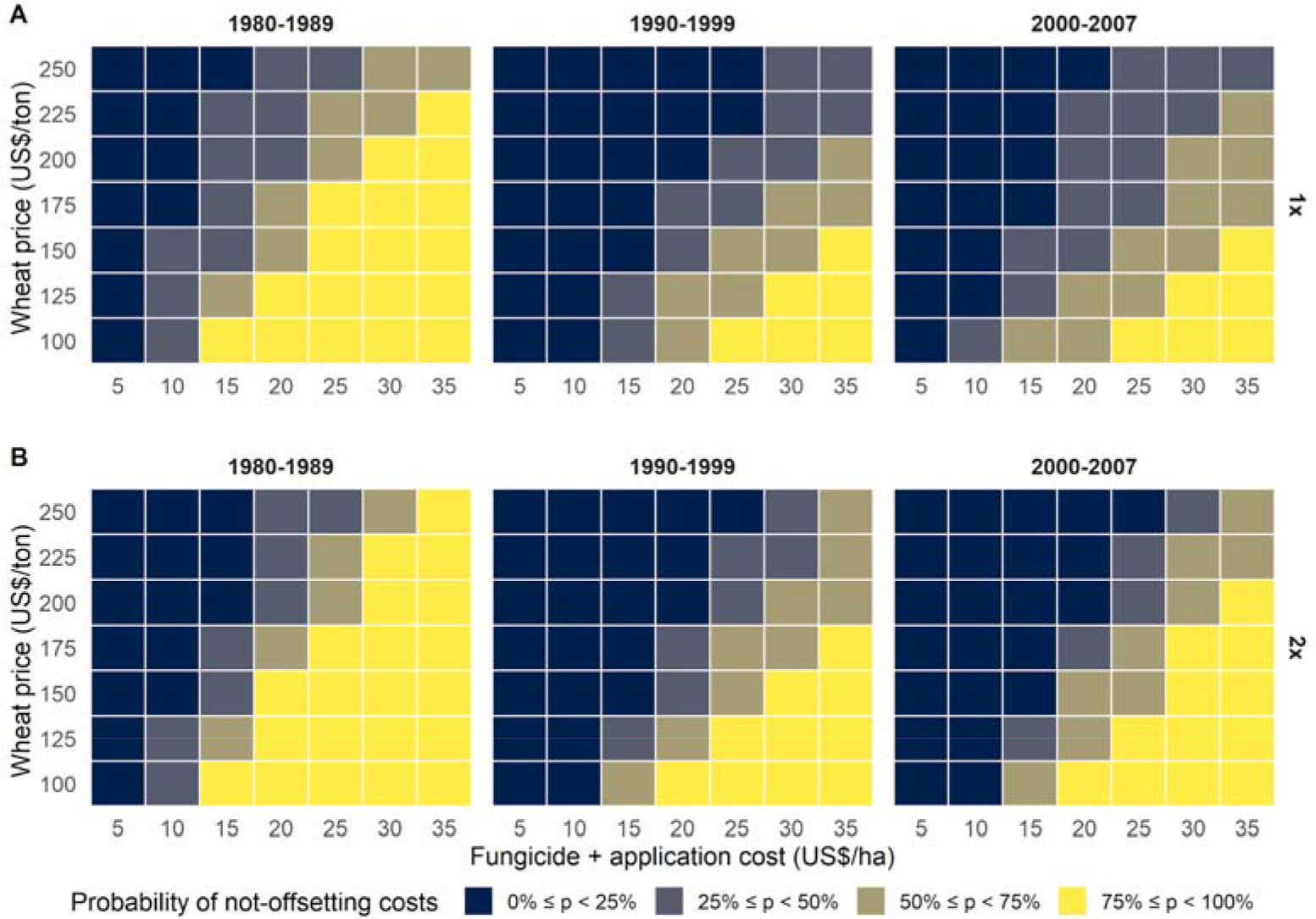
Probability categories of non-offsetting on fungicide investment for different scenarios of wheat prices and fungicide costs (product price + operational costs) for **A**, one or **B**, two sprays (first spray at early to mid-flowering and a second 7 to 10 days later) for Fusarium head blight (FHB) control. Probability for each time period, encompassing the periods prior (1980-1989) and after (1990-1999, 2000-2007) FHB resurgence, was calculated using the estimates of the mean difference 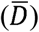, and respective between-year standard deviation (*σ^2^*) obtained from the yield estimates with and without fungicide application for 28 growing seasons in Brazil.

**Figure 5.**
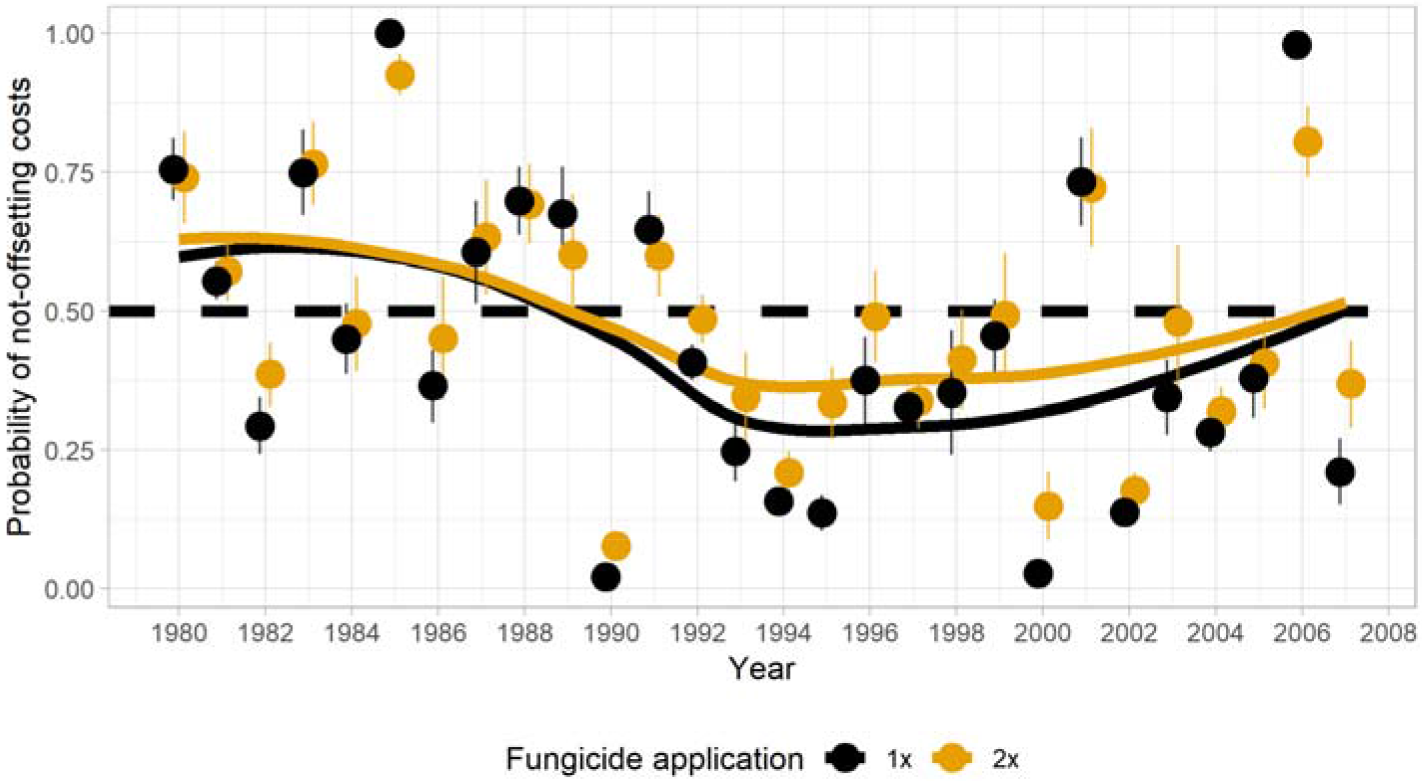
Temporal series of the mean probability, and respective 95% confidence interval, of non-offsetting the costs (probability of loss) for benefit-cost ratios (wheat price / fungicide cost in US$, see ranges in Figure 4) ranging from 5 to 15. The probabilities were calculated for both one and two fungicide application

## Discussion

Using a meta-analytic approach, our quantitative information confirms that FHB is an important yield-limiting disease of spring wheat in the subtropics (Casa et al. 2004; Panisson et al 2003; Spolti et al. 2015). This study also provides numerical estimates of yield loss trends due to FHB in a temporal series of 28-years by linking the meta-analytical estimate parameters to a crop and a disease model to estimate yield losses. Our estimates of yield loss due to FHB followed trend similar to that reported by Reis (1996) and Reis and Carmona (2013) using a completely different approach; those studies estimated 5.4% of yield losses spanning 10 years (1984-1994) and 21.6% for the 2000 to 2010 period, respectively. This suggests that our estimates match quite well with the magnitude and trend of increase in yield loss verified in Brazil after FHB resurgence, with no major trend changes afterward.

Our results highlight that baseline estimates of yield (population-average intercept) was similar to the estimates for the same wheat type (spring) in the United States (Madden and Paul 2009). Nonetheless, slope estimates differed among these different studies, likely due to their inherent characteristics of the production area. The variance in the parameter estimates of meta-regression models may be reduced by inclusion of additional covariates in the model (Lehner et al. 2017; Madden and Paul 2009). Although baseline yield did not affect the population-average slope, intercept values differed between trials representing a low or a high-yielding situation, which was not an unexpected result. A similar pattern was found in the US study (Madden and Paul 2009) where population-average slopes did not differ between spring and winter wheat, which typically represent low and high-yielding wheat, respectively. Given that intercept values varied among the studies, the magnitude of the damage coefficient, which depends on both coefficients, differed between the groups within each country, although there were some similarities between countries for some of the conditions. For example, for a high-yielding spring wheat situation, the *Dc* calculated in our study (1.05%^−1^) was very close to an overall (unconditioned) *Dc* calculated for spring wheat in the United States (0.99%^−1^) (Madden and Paul 2009), which indicates that spring wheat yields in the US were higher than in Brazil on average. In a further US study with data for soft red winter wheat, the damage coefficient due to FHB was slightly greater (1.17%^−1^) and neither wheat cultivar nor the presence of foliar disease Stagonospora leaf blotch (SLB) significantly affected the slopes (Salgado et al 2015).

The increased damage coefficient for subtropical spring wheat in the low-yielding situation may be due to the additional effect of unobserved biotic and abiotic stresses that weakened the plant and its defenses against FHB (Al-Khatib and Paulsen 1999; Asseng et al. 2015; Buck et al. 2007; Sharma et al 2007). In addition, we cannot rule out, given we used data from fungicide trials, the potential effect of fungicides, especially DMI + QoI mixtures applied during flowering, controlling flag-leaf diseases like tan spot in Brazil. This can affect the damage coefficients due to reduced impact of foliar disease, thus protecting yield (Blandino et al 2006, 2011; Wegulo et al. 2011).

Using a model-based approach that linked a weather-based FHB model with a wheat simulation model and the meta-analytical estimate of the damage coefficient, FHB-induced yield loss trends were quantified at magnitudes that matched earlier literature reports. For example, a quantitative review of yield losses, measured using a destructive sampling approach in fungicide trials spanning 21 years (1984-2010) in southern Brazil, reported average losses due to FHB shifted from 5.4% during the period 1984 to 1994 to 21.6% for the period 2000 to 2010 (Reis et al. 1996; Reis and Carmona 2013). Our simulations confirmed this shift by showing that FHB changed from a secondary disease to an economically damaging disease of wheat after the 1990s. Several reasons explain the re-emergence of FHB, which has been reported in several other regions worldwide, including the wider adoption of conservation tillage practices and increase of wheat-corn rotations (Dill-Macky and Jones 2000; Schaafsma et al. 2001; McMullen et al 1997).

In spite of this knowledge, the effectiveness of cultural practices for managing FHB in Brazil was previously question (Reis 1990) and current evidence that supports the claim in subtropical cropping conditions includes: a) overwintering of several grasses harbor the pathogen (Reis 1990); b) presence of airborne inoculum throughout the year (Reis 1988; Fernandes, 1997); and c) non-influence of the presence of within-field maize residues on FHB risk (Spolti et al. 2015). Collectively, results from our simulations based only on seasonal weather and not taking into account changes in agronomic practices (that promote inoculum build-up), confirm that the re-emergence of FHB in Brazil has been more likely due to variation in seasonal weather patterns, as previously suggested (Del Ponte et al. 2009) rather than cultural practices (Fernandes 1997). Our results are important for guiding priorities for management, including breeding efforts to mitigate both direct and indirect losses due to FHB, especially given the increased production costs from reliance on fungicides recommended for protecting yield and reducing mycotoxin contamination (McMullen et al. 2012; Wegulo et al. 2015).

Prior to the early 1990s, the profitability of a single fungicide spray during flowering for FHB control was questioned due the sporadic nature and mostly mild FHB epidemics in the preceding decades (Picinini and Fernandes 1998; Reis et al. 1996). However, accurate yield loss estimates, using a destructive sampling approach during the early 2000s, contradicted the previous recommendation and showed economic benefits from using one fungicide spray at full flowering (Panisson et al 2003). Our model-based estimates of yield losses and fungicide profitabilities showed that the risk of not-offsetting a single fungicide sprays was predominantly high prior to the resurgence (< 1990s), evidenced by the low frequency of more profitable conditions, and agreeing with current recommendation for the period (Reis et al. 1996). The profitability tended to increase in general after the 1990s, when a single fungicide spray, but not often two sprays, was more likely to pay-off. Caution is needed in interpretation because we are not taking into account the effect of fungicide sprays on other traits affected by FHB such as test weight and mycotoxin, which downgrade grain value (Salgado et al 2014, 2015).

We found that the yield loss time series was not affected by planting period, although it is expected that there may be annual variation since previous reports indicated increasing risk of FHB for later planting dates (after June 15) compared to prior dates (Lima et al. 2005, 2006). Therefore, it is difficult to ascertain any general recommendation on planting dates for managing FHB, though it is important mentioning that our disease model does not take an increase in airborne inoculum levels into account, which tends to peak later in the season (Del Ponte et al. 2009). Given the dependence of FHB epidemics on weather conditions during a relatively short period (flowering) (Del Ponte et al. 2005), diversification of planting dates has been proposed as a way to minimize risk through an escape mechanism (Reis and Carmona, 2013). Our simulations indeed showed significant within-year variations in yield losses, varying in magnitude across the years.

Our approach for modeling yield loss may be useful for risk assessment and design of long-lasting, yet profitable, contingency tactics to management FHB in wheat. Under the current scenario, one fungicide spray (tebuconazole) during flowering is more likely a profitable decision than applying two sprays, for which there is greater uncertainty. The damage coefficients can be useful in decision support tools like apps embed with real time predictions of disease severity and economic scenarios (Willbur et al. 2018). Situations at which an additional fungicide spray is worth the cost need to be better explored under scenarios of different cultivars and fungicide chemistry on other quality parameters that downgrade grain such as test weight and mycotoxin contamination.

## Acknowledgements

The authors thank to Dr. A. Lazzaretti and Dr. JM Fernandes for providing simulated yield data from the AgroWEB system. Research conducted by Paul Esker was also supported by the USDA National Institute of Food and Federal Appropriations under Project PEN04660 and Accession number 1016474. First and Last author are thankful to CAPES and CNPq for the scholarship and research fellow support, respectively.

## Literature cited

Asseng, S., Ewert, F., Martre, P., Rötter, R. P., Lobell, D. B., Cammarano, D., et al. 2015. Rising temperatures reduce global wheat production. Nat. Clim. Change. 5:143–147.

Al-Khatib, K., and Paulsen, G. M. 1999. High-temperature effects on photosynthetic processes in temperate and tropical cereals. Crop Sci. 39:119.

Blandino, M., Minelli, L., and Reyneri, A. 2006. Strategies for the chemical control of Fusarium head blight: Effect on yield, alveographic parameters and deoxynivalenol contamination in winter wheat grain. Eur. J. Agron. 25:193–201.

Blandino, M., Pascale, M., Haidukowski, M., and Reyneri, A. 2011. Influence of agronomic conditions on the efficacy of different fungicides applied to wheat at heading: effect on flag leaf senescence, Fusarium head blight attack, grain yield and deoxynivalenol contamination. Ital. J. Agron. 6:32.

Brum, A. L.; Heck, C. R. 2005. A economia do trigo no Rio Grande do Sul: breve histórico do cereal na economia do estado. Revista Análise. 16:29–44.

Buck, H. T., Nisi, J. E., Salomón, N. 2007. Wheat production in stressed environments. Springer, Dordrecht, NL.

Casa, R. T., Bogo, A., Moreira, É. N., and Kuhnem Junior, P. R. 2007. Época de aplicação e desempenho de fungicidas no controle da giberela em trigo. Ciênc. Rural. 37:1558–1563.

Casa, R. T., Reis, E. M., Blum, M. M. C., Bogo, A., Scheer, O., and Zanata, T. 2004. Danos causados pela infecção de Gibberella zeae em trigo. Fitopatol. Bras. 29:289–293.

CEPEA. 2018. Centro de Estudos Avançados em Economia Aplicada. Preço médio do trigo CEPEA/ESALQ. Piracicaba, SP. https://www.cepea.esalq.usp.br/br/indicador/trigo.aspx.

CONAB, 2018. Companhia Nacional de Abastecimento. Acompanhamento de safra brasileira de grão. Brasília, DF. https://www.conab.gov.br/info-agro/safras

Dalla Lana, F., Ziegelmann, P. K., Maia, A. de H. N., Godoy, C. V., and Del Ponte, E. M. 2015. Meta-analysis of the relationship between crop yield and soybean rust severity. Phytopathology. 105:307–315.

Del Ponte, E. M., Fernandes, J. M. C., Pavan, W., and Baethgen, W. E. 2009. A model-based assessment of the impacts of climate variability on Fusarium head blight seasonal risk in southern Brazil. J. Phytopathol. 157:675–681.

Del Ponte, E. M., Fernandes, J. M. C., and Pierobom, C. R. 2005. Factors affecting density of airborne Gibberella zeae inoculum. Fitopatol. Bras. 30:55–60.

Del Ponte, E. M., Spolti, P., Ward, T. J., Gomes, L. B., Nicolli, C. P., Kuhnem, P. R., et al. 2015. Regional and field-specific factors affect the composition of Fusarium head blight pathogens in subtropical no-till wheat agroecosystem of Brazil. Phytopathology. 105:246–254.

Deuner, C. C. 2009. Eficácia agronômica de fungicidas no controle de giberela na cultura do trigo. Pages 269–274 in: Controle de doenças em plantas. FUNDACEP 1993-2008. Resultados de pesquisa. Acervo histórico. Divulgação técnica N° 3. Cruz Alta.

Dill-Macky, R., and Jones, R. K. 2000. The effect of previous crop residues and tillage on Fusarium head blight of wheat. Plant Dis. 84:71–76.

Edwards Molina, J. P., Paul, P. A., Amorim, L., da Silva, L. H. C. P., Siqueri, F. V., Borges, E. P., et al. 2018. Meta-analysis of fungicide efficacy on soybean target spot and cost-benefit assessment. Plant Pathol. 68:94–106.

Edwards, S. G., and Godley, N. P. 2010. Reduction of Fusarium head blight and deoxynivalenol in wheat with early fungicide applications of prothioconazole. Food Addit. Contam. Part A. 27:629–635.

Feksa, H. R., Gardiano, C. G., Duhatschek, B., Proença, C., and Tessmann, D. J. 2014a. Ensaio manejo químico de giberela na cultura do trigo cultivar BRS Guamirim. In: Proceedings of Reunião da comissão brasileira de pesquisa de trigo e triticale (CD-ROM).

Feksa, H. R., Gardiano, C. G., Duhatschek, B., Proença, C., and Tessmann, D. J. 2014b. Ensaio manejo químico de giberela na cultura do trigo cultivar CD 105. In: Proceedings of Reunião da comissão brasileira de pesquisa de trigo e triticale (CD-ROM). Embrapa Trigo, Passo Fundo, Brazil.

Feldmann, N.A., Mühl, F.R., Hahn, L., Hoffmann, J.T.R., Klein, R., Zambiazi, M.P., Martini, A., Jantsch, M., Lima, A., 2014. Avaliação de fungicidas no controle de giberela do trigo. Anais eletrônicos da VIII Reunião da Comissão Brasileira de Pesquisa de Trigo e Triticale. Canela, 2014. CD - ROM.

Fernandes, J. M. C. 1997. As doenças das plantas e o sistema de plantio direto. Revisão Anual de Patologia de Plantas. 5:317–352.

Goswami, R. S., and Kistler, H. C. 2004. Heading for disaster: Fusarium graminearum on cereal crops. Mol. Plant Pathol. 5:515–525.

Guterres, C. W., Bruinsma, J. da S., Seidel, G. 2014. Efficacy of fungicides for FHB control and reduction of deoxynivalenol in wheat. In: Proceedings of 5th International symposium on Fusarium head blight. Florianópolis, 2015. CD-ROM.

Kazan, K., Gardiner, D. M., and Manners, J. M. 2012. On the trail of a cereal killer: recent advances in Fusarium graminearum pathogenomics and host resistance. Mol. Plant Pathol. 13:399–413.

Kikot, G. E., Moschini, R., Consolo, V. F., Rojo, R., Salerno, G., Hours, R. A., et al. 2011. Occurrence of different species of Fusarium from wheat in relation to disease levels predicted by a weather-based model in Argentina pampas region. Mycopathologia. 171:139–149.

Landschoot, S., Audenaert, K., Waegeman, W., De Baets, B., and Haesaert, G. 2013. Influence of maize–wheat rotation systems on Fusarium head blight infection and deoxynivalenol content in wheat under low versus high disease pressure. Crop Prot. 52:14–21.

Lazzaretti, A. T., Fernandes, J. M., and Pavan, W. 2015. Calibração do cropsim-wheat para simulação do desenvolvimento e rendimento de grão de trigo no Sul do Brasil. Rev. Bras. Ciênc. Agrár. - Braz. J. Agric. Sci. 10:356–364.

Lehner, M. S., Pethybridge, S. J., Meyer, M. C., and Del Ponte, E. M. 2017. Meta-analytic modelling of the incidence-yield and incidence-sclerotial production relationships in soybean white mould epidemics. Plant Pathol. 66:460–468

Lima, M. I. P. M., Só e Silva, M., Schereen, P. L., Del Duca, I. J. A., Pires, J. L., Nascimento, A., Jr. 2005. Avaliação de giberela em genótipos de trigo do ensaio estadual de cultivares, na região de Passo Fundo, em 2004. Documento online 52. Passo Fundo, Embrapa Trigo, 11p. http://www.cnpt.embrapa.br/biblio/do/p_do52.htm

Lima, M. I. P. M., Só e Silva, M.; Caierão, E., Schereen, P. L., Del Duca, I. J. A., Pires, J. L., 2006. Avaliação de giberela em genótipos de trigo do ensaio estadual de cultivares, na região de Passo Fundo, em 2005. Documento online 66. Passo Fundo, Embrapa Trigo, 7p. http://www.cnpt.embrapa.br/biblio/do/p_do66.htm.

Machado, F. J., Santana, F. M., Lau, D., and Del Ponte, E. M. 2017. Quantitative review of the effects of triazole and benzimidazole fungicides on Fusarium head blight and wheat yield in Brazil. Plant Dis. 101:1633–1641.

Madden, L. V., and Paul, P. A. 2009. Assessing heterogeneity in the relationship between wheat yield and Fusarium head blight intensity using random-coefficient mixed models. Phytopathology. 99:850–860.

Madden, L. V., Piepho, H.-P., and Paul, P. A. 2016. Statistical models and methods for network meta-analysis. Phytopathology. 106:792–806.

Maffini, F. S., Arbugeri, F. E., Canova, E., Dahmer, J., Pinto, F. F., Ebone, A., Uebel, J. D., Balardin, R. S. 2012. Programas de controle químico de giberela (Fusarium graminearum) na cultura do trigo. In: XVI Simpósio de Ensino, Pesquisa e Extensão. 2012. (CD-ROM).

McMullen, M., Bergstrom, G., De Wolf, E., Dill-Macky, R., Hershman, D., Shaner, G., et al. 2012. A unified effort to fight an enemy of wheat and barley: Fusarium head blight. Plant Dis. 96:1712–1728.

McMullen, M., Jones, R., and Gallenberg, D. 1997. Scab of wheat and barley: a re-emerging disease of devastating impact. Plant Dis. 81:1340–1348.

Moschini, R. C., and Fortugno, C. 1996. Predicting wheat head blight incidence using models based on meteorological factors in Pergamino, Argentina. Eur. J. Plant Pathol. 102:211–218.

Panisson, E., Reis, E. M., and Boller, W. 2002. Efeito da época, do número de aplicações e de doses de fungicida no controle da giberela em trigo. Fitopatol. Bras. 27:489–494.

Panisson, E., Reis, E. M., and Boller, W. 2003. Quantificação de danos causados pela giberela em cereais de inverno, na safra 2000, em Passo Fundo, RS. Fitopatol. Bras. 28:189–192.

Paul, P. A., Lipps, P. E., Hershman, D. E., McMullen, M. P., Draper, M. A., and Madden, L. V. 2008. Efficacy of triazole-based fungicides for Fusarium head blight and deoxynivalenol control in wheat: a multivariate meta-analysis. Phytopathology. 98:999–1011.

Paul, P. A., Lipps, P. E., and Madden, L. V. 2005. Relationship between visual estimates of Fusarium head blight intensity and deoxynivalenol accumulation in harvested wheat grain: a meta-analysis. Phytopathology. 95:1225–1236.

Paul, P. A., Lipps, P. E., Hershman, D. E., McMullen, M. P., Draper, M. A., and Madden, L. V. 2007. A quantitative review of tebuconazole effect on Fusarium head blight and deoxynivalenol content in wheat. Phytopathology 97:211–220.

Paul, P. A., Lipps, P. E., Hershman, D. E., McMullen, M. P., Draper, M. A., and Madden, L. V. 2008. Efficacy of triazole-based fungicides for Fusarium head blight and deoxynivalenol control in wheat: A multivariate meta-analysis. Phytopathology 98:999–1011.

Paul, P. A., Madden, L. V., Bradley, C. A., Robertson, A. E., Munkvold, G. P., Shaner, G., et al. 2011. Meta-analysis of yield response of hybrid field corn to foliar fungicides in the U.S. Corn Belt. Phytopathology. 101:1122–1132.

Picinini, E. C., Fernandes, J. M. C. 1998. Avaliação de fungicidas no controle de giberela em trigo. Fitopatol Bras. 23:270.

Pizolotto, C. A., and Boller, W. 2015. Spray nozzles and frequency of fungicides applications to the control of Fusarium head blight in wheat. In: Proceedings of 5th International symposium on Fusarium head blight. Florianópolis, 2015. CD-ROM.

R Core Team. 2013. R: A language and environment for statistical computing. R Foundation for Statistical Computing, Vienna, Austria. URL http://www.R-project.org/.

Reis, E. M. 1988. Quantificação de propágulos de Gibberella zeae no ar através de armadilhas de esporos. Fitopatol Bras 13:324–327.

Reis E. M. 1990. Perithecial formation of Gibberella zeae on senescent stems of grasses under natural conditions. Fitopatol Bras 15:52–54.

Reis, E. M., and Carmona, M. A. 2013. Integrated disease management of Fusarium head blight. Pages 159–173 in: Fusarium head blight in latin America. Magliano, T. M. A., Chulze, S. N, eds. Springer, Dordrecht, NL.

Reis, E. M., Blum, M. M. C., Casa, R., Medeiros, C. A., 1996. Grain losses caused by the infection of wheat heads by Gibberella zeae in southern Brazil, from 1984 to 1994. Summa Phytopathol. 22:134–137.

Salgado, J. D., Madden, L. V., and Paul, P. A. 2014. Efficacy and economics of integrating in-field and harvesting strategies to manage Fusarium head blight of wheat. Plant Dis. 98:1407–1421.

Salgado, J. D., Madden, L. V., and Paul, P. A. 2015. Quantifying the effects of Fusarium head blight on grain yield and test weight in soft red winter wheat. Phytopathology. 105:295–306.

Salgado, J. D., Wallhead, M., Madden, L. V., and Paul, P. A. 2011. Grain harvesting strategies to minimize grain quality losses due to Fusarium head blight in wheat. Plant Dis. 95:1448–1457.

Santana, F. M., Lau, D., Aguilera, J. G., Sbalcheiro, C. C., Feksa, H., Floss, L. G., and Guterres, C. W. 2016a. Eficiência de fungicidas para controle de Gibberella zeae em trigo: resultados dos Ensaios Cooperativos - Safra 2013. Comunicado Técnico 362. Embrapa Trigo, Passo Fundo, Brazil.

Santana, F. M., Lau, D., Cargnin, A., Seixas, C. D. S., Schipanski, C. A., Feksa, H. R., Wesp, C., Blum, M., and Bassoi, M. C. 2014. Eficiência de fungicidas para controle de giberela em trigo: resultados dos ensaios cooperativos - safra 2012. Comunicado Técnico 336. Embrapa Trigo, Passo Fundo, Brazil.

Santana, F. M., Lau, D., Maciel, J. L. N., Cargnin, A., Seixas, C. D. S., Bassoi, M. C., Schipanski, C. A., Feksa, H., Casa, R. T., Wesp, C., Navarini, L., and Blum, M. 2012. Eficiência de fungicidas para controle de giberela em trigo: resultados dos ensaios cooperativos - safra 2011. Comunicado Técnico 23. Embrapa Trigo, Passo Fundo, Brazil.

Santana, F. M., Lau, D., Sbalcheiro, C. C., Feksa, H., Guterres, C. W., and Venâncio, W. S. 2016b. Eficiência de fungicidas para controle de Gibberella zeae em trigo: resultados dos Ensaios Cooperativos - Safra 2015. Comunicado Técnico 368. Embrapa Trigo, Passo Fundo, Brazil.

Santana, F. M., Lau, D., Sbalcheiro, C. C., Schipanski, C. A., Seixas, C. D. S., Feksa, H., Floss, L. G., Guterres, C. W., and Venâncio, W. S. 2016c. Eficiência de fungicidas para controle de Gibberella zeae em trigo: resultados dos Ensaios Cooperativos - Safra 2014. Comunicado Técnico 364. Embrapa Trigo, Passo Fundo, Brazil.

Savary, S., Nelson, A. D., Djurle, A., Esker, P. D., Sparks, A., Amorim, L., et al. 2018. Concepts, approaches, and avenues for modelling crop health and crop losses. Eur. J. Agron. 100:4–18.

Schaafsma, A. W., Ilinic, L. T.-, Miller, J. D., and Hooker, D. C. 2001. Agronomic considerations for reducing deoxynivalenol in wheat grain. Can. J. Plant Pathol. 23:279–285.

Schaafsma, A. W., Tamburic-Ilincic, L., and Hooker, D. C. 2005. Effect of previous crop, tillage, field size, adjacent crop, and sampling direction on airborne propagules of Gibberella zeae/Fusarium graminearum, Fusarium head blight severity, and deoxynivalenol accumulation in winter wheat. Can. J. Plant Pathol. 27:217–224.

Shah, D. A., Molineros, J. E., Paul, P. A., Willyerd, K. T., Madden, L. V., and De Wolf, E. D. 2013. Predicting Fusarium head blight epidemics with weather-driven pre- and post-anthesis logistic regression models. Phytopathology. 103:906–919.

Shah, L., Ali, A., Yahya, M., Zhu, Y., Wang, S., Si, H., et al. 2018. Integrated control of Fusarium head blight and deoxynivalenol mycotoxin in wheat. Plant Pathol. 67:532–548.

Sharma, R. C., Duveiller, E., and Ortiz-Ferrara, G. 2007. Progress and challenge towards reducing wheat spot blotch threat in the Eastern Gangetic Plains of South Asia: Is climate change already taking its toll? Field Crops Res. 103:109–118.

Spolti, P., Del Ponte, E. M., Dong, Y., Cummings, J. A., and Bergstrom, G. C. 2014. Triazole sensitivity in a contemporary population of Fusarium graminearum from New York wheat and competitiveness of a tebuconazole-resistant isolate. Plant Dis. 98:607–613.

Spolti, P., Guerra, D. S., Badiale-Furlong, E., and Del Ponte, E. M. 2013. Single and sequential applications of metconazole alone or in mixture with pyraclostrobin to improve Fusarium head blight control and wheat yield in Brazil. Trop. Plant Pathol. 38:85–96.

Spolti, P., Shah, D. A., Fernandes, J. M. C., Bergstrom, G. C., and Del Ponte, E. M. 2015. Disease risk, spatial patterns, and incidence-severity relationships of Fusarium head blight in no-till spring wheat following maize or soybean. Plant Dis. 99:1360–1366.

Venancio, W.S., Santos, T., Boratto, V.N.M., Skodowski, L.H. 2014. Manejo de doenças do trigo com fungicidas, alternando a mistura pronta de isoftalonitrila + triazol. Anais eletrônicos da VIII Reunião da Comissão Brasileira de Pesquisa de Trigo e Triticale. Canela, 2014. CD-ROM.

Viana, C., 2013. Giberela em trigo: sobrevivência, reação de cultivares e controle químico. 111 f. Dissertação (Mestrado em Agronomia) – Faculdade de Agronomia e Medicina Veterinária da Universidade de Passo Fundo, Passo Fundo, 2013.

Wegulo, S. N., Baenziger, P. S., Hernandez Nopsa, J., Bockus, W. W., and Hallen-Adams, H. 2015. Management of Fusarium head blight of wheat and barley. Crop Prot. 73:100–107.

Wegulo, S. N., Bockus, W. W., Nopsa, J. H., De Wolf, E. D., Eskridge, K. M., Peiris, K. H. S., et al. 2011. Effects of integrating cultivar resistance and fungicide application on Fusarium head blight and deoxynivalenol in winter wheat. Plant Dis. 95:554–560.

Willbur, J. F., Fall, M. L., Byrne, A. M., Chapman, S. A., McCaghey, M. M., Mueller, B. D., et al. 2018. Validating Sclerotinia sclerotiorum apothecial models to predict Sclerotinia stem rot in soybean (Glycine max) fields. Plant Dis. 102:2592–2601.

Willyerd, K. T., Li, C., Madden, L. V., Bradley, C. A., Bergstrom, G. C., Sweets, L. E., et al. 2012. Efficacy and stability of integrating fungicide and cultivar resistance to manage Fusarium head blight and deoxynivalenol in wheat. Plant Dis. 96:957–967.

Wood, S. N. 2017. Generalized additive models: an introduction with R. Chapman and

